# Periodical cicadas suffer legacy effects of century-old forest removal

**DOI:** 10.1101/2025.11.24.690214

**Authors:** Ian Chan, Guerin E. Brown, Fabiola Castaneda-Santiago, Jacky S. Chitty, Tamra A. Elliott, Mattie J. Gose, Charlotte M. Hanfland, Stephen D. Hendrix, Carter R. Leerhoff, Susan K. Meerdink, Adam M. Skibbe, MaKella J. Steffensen, Christian L. Weinrich, Andrew A. Forbes

## Abstract

Periodical cicada (*Magicicada* spp.) ranges have continued to decline even as their forest habitat has rebounded from historic lows. This counterintuitive trajectory may be a consequence of *Magicicada*’s limited dispersal capabilities combined with their requirement that individual host trees survive across decades. We studied the relationship between >150 years of landscape change and brood XIII periodical cicada activity in Johnson County, IA. We found that the presence of *Magicicada* breeding populations was better predicted by tree cover in 1930 than by present-day tree cover, with cicadas primarily relegated to sites that did not fall below 17.4% tree cover. While total forest has increased since 1930, this is a net increase, inclusive of considerable turnover every 17 years, and cicada ranges have declined more than they have expanded. A model based on empirical measures of forest turnover and cicada dispersal distances suggests that cicada ranges will continue to decline under nearly all conditions. A managed relocation of 20,126 adult *Magicicada* to a restored forest failed, reinforcing that dispersal to new sites cannot be artificially accelerated to offset declines. Preservation of forest habitat across centuries may be the only bollard standing between periodical cicadas and their extinction.

## Text

Periodical cicadas (genus *Magicicada*) are endemic North American insects with adults that emerge synchronously and *en masse* in 13- or 17-year intervals (Kritsky, 2004). The large densities of individuals that emerge within just a few weeks in early summer (>1 million per acre at some sites; Dybas & Dybas, 1962) satiate local predators, leaving millions to survive, mate, and reproduce (Karban 1982; Williams et al., 1993). During emergence years they provide an important food resource to small mammals and birds (Koenig et al., 2005; Marcello et al., 2008), directly increase nitrogen availability in soils (Yang 2004; Setälä et al., 2022) and indirectly benefit diversity and abundance of other forest insects (Getman-Pickering et al., 2023). Their large body sizes, extraordinarily high densities, and loud (50-80 dB), diurnal chorusing also attract considerable public and media attention (Simon et al., 2022).

Despite their massive numbers, periodical cicadas appear to be experiencing a general decline. Decades-long mapping projects of both 13- and 17-year *Magicicada* broods (“brood” = a group of cicadas, often including two or more different species) show ranges shrinking over time (Cooley et al., 2004, 2013, 2016; Gilbert & Klass 2006) and at least two broods (XI and XXI) have become extinct (Cooley et al., 2013; Marlatt 1907; Manter 1974). The situation for periodical cicadas thus has a timbre reminiscent of other North American fauna like the passenger pigeon, Carolina parakeet, and Rocky Mountain locust – animals of extravagant abundance that nevertheless declined towards eventual extinction (Gaston & Fuller, 2008).

Loss of forests in the Eastern USA on a regional scale (Thompson et al. 2013) is one factor connected to periodical cicada range reduction (Cooley et al. 2004; Gilbert & Klass 2006), largely because the entire life cycle of *Magicicada* depends on deciduous trees. Emerging adults feed on tree xylem fluids, male *Magicicada* chorus in tree canopies to attract females, and females lay eggs into terminal branches of trees (Williams & Simon, 1995; Reiter et. al., 2023). Most importantly, nymphal stages feed on tree root xylem fluids for their entire 13- or 17-year development period such that if a host tree dies so do the associated nymphs underground (Perkovich & Ward 2023). Thus, clearcutting a forest would remove all cicadas present at that site.

While absolute tree cover has increased in the last half century in regions where periodical cicadas are endemic, the ranges of *Magicicada* have not increased correspondingly (Gilbert & Klass 2006; Jungst et al., 1998) and some forests undisturbed for > 50 years have had lower *Magicicada* densities than in previous censuses (Maier 1982). This incongruence between increased availability of acceptable habitat and continuing *Magicicada* range contractions may reflect a sensitivity to historical land use change peculiar to periodical cicadas. Specifically, not only are *Magicicada* easily lost from a site if their host trees are removed, but they may also have difficulty recolonizing reforested sites – even after a century or more – because of their intermittent opportunities for dispersal (just 5*–*7 times per century) and short dispersal distances (generally <50m; Karban 1981). If this hypothesis is correct, historical forest cover should have a stronger influence than contemporary forest cover on current cicada distributions.

### Historical forest removal reduced Magicicada ranges

Johnson County, Iowa, USA (1610 km^2^) is ideal for measuring the effect of land use on *Magicicada* distributions because it has a well-documented history of land use change (including survey data and aerial photography) as well as historical and contemporary records of cicadas. Johnson County is part of the ancestral range of Brood XIII 17-year cicadas (Marlatt 1907) and a formal survey of the county from 1836-1859 documented large continuous swaths of land (∼465.3 km^2^; 28.9% of the county) defined either as “timber”, “oak barrens”, or “grove”, indicative of tree cover acceptable to *Magicicada* (Figure S1). We used aerial photos to classify tree cover in 1930 and 2023 and scoured Iowa City area newspapers for all mention of periodical cicada activity (Supplemental Methods). Newspapers from 1871, 1888, and 1905 describe mass cicada emergences within the limits of today’s Iowa City, where historical tree cover had extended (Table S1). By 1930, encroaching farmland, pastureland, and urbanization had reduced tree cover to just 181.1 km^2^ (11.2% of the county; Figure S2). However, articles from post-1905 that mention periodical cicadas are either purely anticipatory or refer exclusively to emergences at sites outside of Iowa City (Table S1), demonstrating that *Magicicada* ranges had retreated. By 2023, tree cover in the county had increased to 206.7 km^2^ (Figure S3) but a 2007 survey showed *Magicicada* limited to a small area of the county (Cooley et al., 2016).

Prior to the 2024 emergence of Brood XIII cicadas in Johnson Co., we surveyed 16 forested sites (Figure 1) for the presence of pre-emergence tunnels, which *Magicicada* dig ∼1 month before exiting soils (Williams & Simon 1995). Of the 16 survey sites, 14 were inside the 1836-1859 range of historically forested areas, including seven sites (#8-#14) in Iowa City. We found evidence of cicada tunnels at 13 sites but also measured considerable within site-variation. Only five sites (#1-#5) had mean densities significantly greater than 0.5 tunnels/m^2^ (Table 1). Only at those same five sites did we see cicadas emerge in large numbers, hear sustained chorusing, observe adults mating, and note tree damage indicative of cicada oviposition (Table 1). These sites were all in a limited area of the county several km north of Iowa City.

**Table 1.** Pre- and post-emergence evidence of periodical cicadas at 16 forested sites in Johnson Co., IA. All sites except for #6 & #15 were in areas classified as having trees during the 1836-1859 GLO survey.

| Site # | Mean ±SE<br>Emergence<br>Tunnels (m<sup>2</sup>) | Adults<br>Present | Chorusing | Egg-<br>laying<br>on<br>Twigs |
| --- | --- | --- | --- | --- |
| 1 | 7.38 ± 1.19 | Yes | Loud | Yes |
| 2 | 2.80 ± 1.14 | Yes | Loud | Yes |
| 3 | 9.80 ± 2.61 | Yes | Loud | Yes |
| 4 | 7.80 ± 2.11 | Yes | Loud | Yes |
| 5 | 5.71 ± 2.03 | Yes | Loud | Yes |
| 6 | 0 | No | No | No |
| 7 | 0.44 ± 0.29 | No | No | No |
| 8 | 0.16 ± 0.13 | No | No | No |
| 9 | 0.02 ± 0.01 | Few<sup>1</sup> | No | No |
| 10 | 0.05 ± 0.05 | No | No<sup>2</sup> | No |
| 11 | 0.05 ± 0.05 | No | No | No |
| 12 | 0.35 ± 0.29 | No | No | No |
| 13 | 1.13 ± 0.94 | Few<sup>1</sup> | No | No |
| 14 | 0 | No | No | No |
| 15 | 0 | No | No | No |
| 16 | 0 | Few<sup>1</sup> | No | No |

<sup>1</sup>While no cicada adults were observed by the authors, either shed nymphal cases (site 9) or photographic evidence on citizen naturalist sites (sites 13 and 16) indicated low-density populations. <sup>2</sup>A single male *M. cassini* call was heard at site 10, but there was no sustained chorusing.

**Fig. 1.**
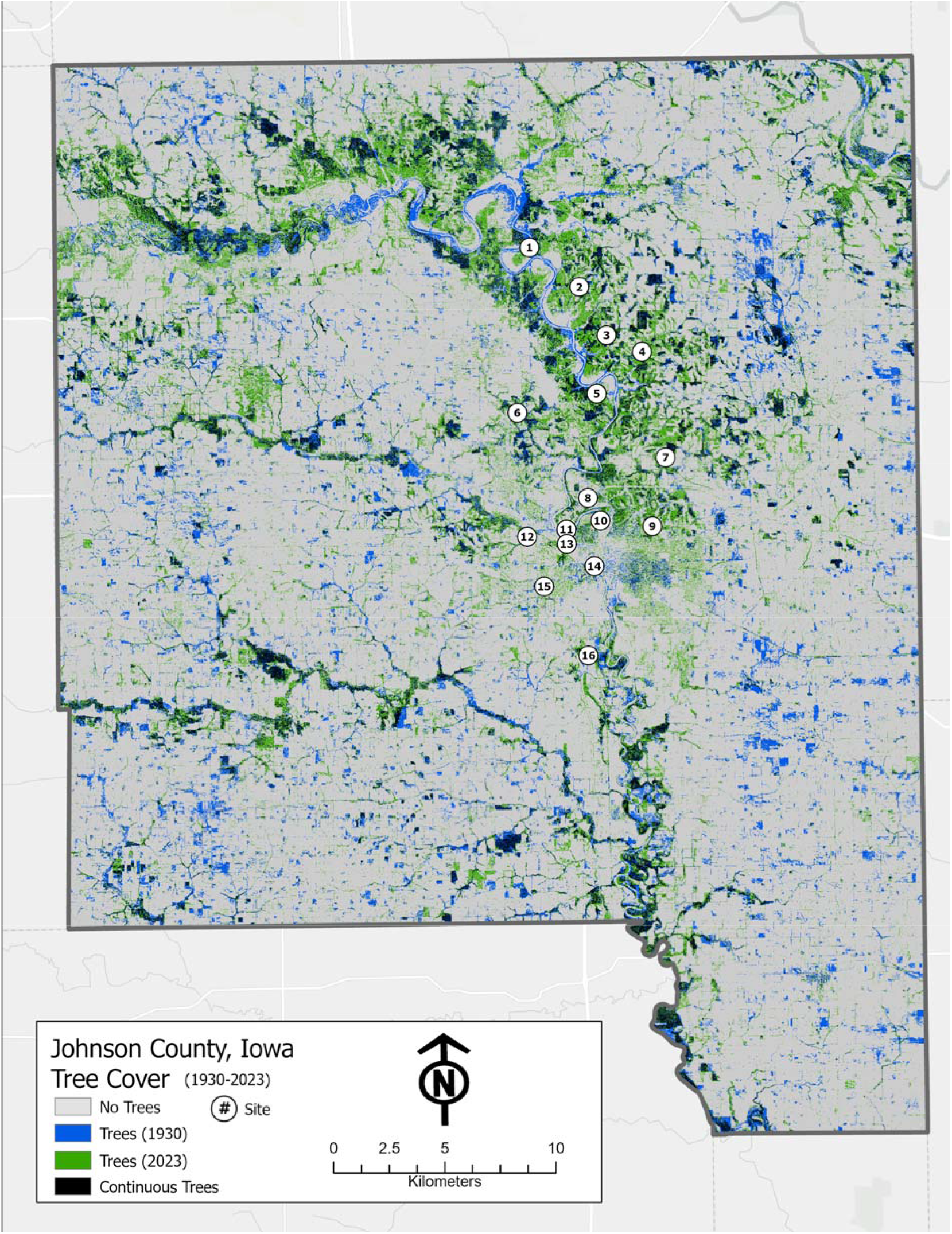
Map of forested landcover in Johnson Co., IA, showing sixteen forested sites surveyed for *Magicicada* before and after their 2024 emergence. Colored pixels represent tree cover. Locations that had trees in 2023 but not in 1930 are shown in green, those with trees only in 1930 are shown in blue, and “continuous” trees (present in both 2023 and 1930) are in black.

### Cicada presence is better predicted by historical forest cover than by contemporary forest cover

After *Magicicada* had emerged across the county, we visited 301 forested sites to listen for male chorusing, a sign of a large and reproducing cicada population (Cooley et al. 2004). We recorded chorusing at 116/301 sites. To determine whether chorusing was best predicted by current or past forest cover, we performed binary logistic regression on chorusing behavior (yes/no) against the percentage of forested landcover surrounding each site in 1836-1859, 1930, and 2023 at seven radii ranging from 0.5 km to 3 km (Supplemental Methods). Percentage forest cover significantly predicted chorusing at all radii tested, for all three timepoints (Figure 2A). Tree cover in 1930 at a 2 km radius provided the best predictive power overall and explained 52.5% of variation (Figure 2A; Table S3). Predictive power of 2023 tree cover was higher than 1836-1859 or 1930 at shorter radii but stayed relatively constant with increasing radius (Figure 2A). This probably reflects only that trees needed to be physically present at the immediate site in 2024 for cicadas to also be present. Tree cover in 1836-1859 also significantly predicted chorusing but power was consistently low (Figure 2A), likely because most forested land had been lost or had turned over (removed and replanted) since 1859. Binomial regression combining three explanatory variables (2023 tree cover at a radius of 0.5 km, 1930 at 3 km, and 1836-1859 at 3km) predicted 64.1% of variation (Table S3), reflecting that the best situation for cicadas is consistent tree cover across time.

**Fig. 2.**
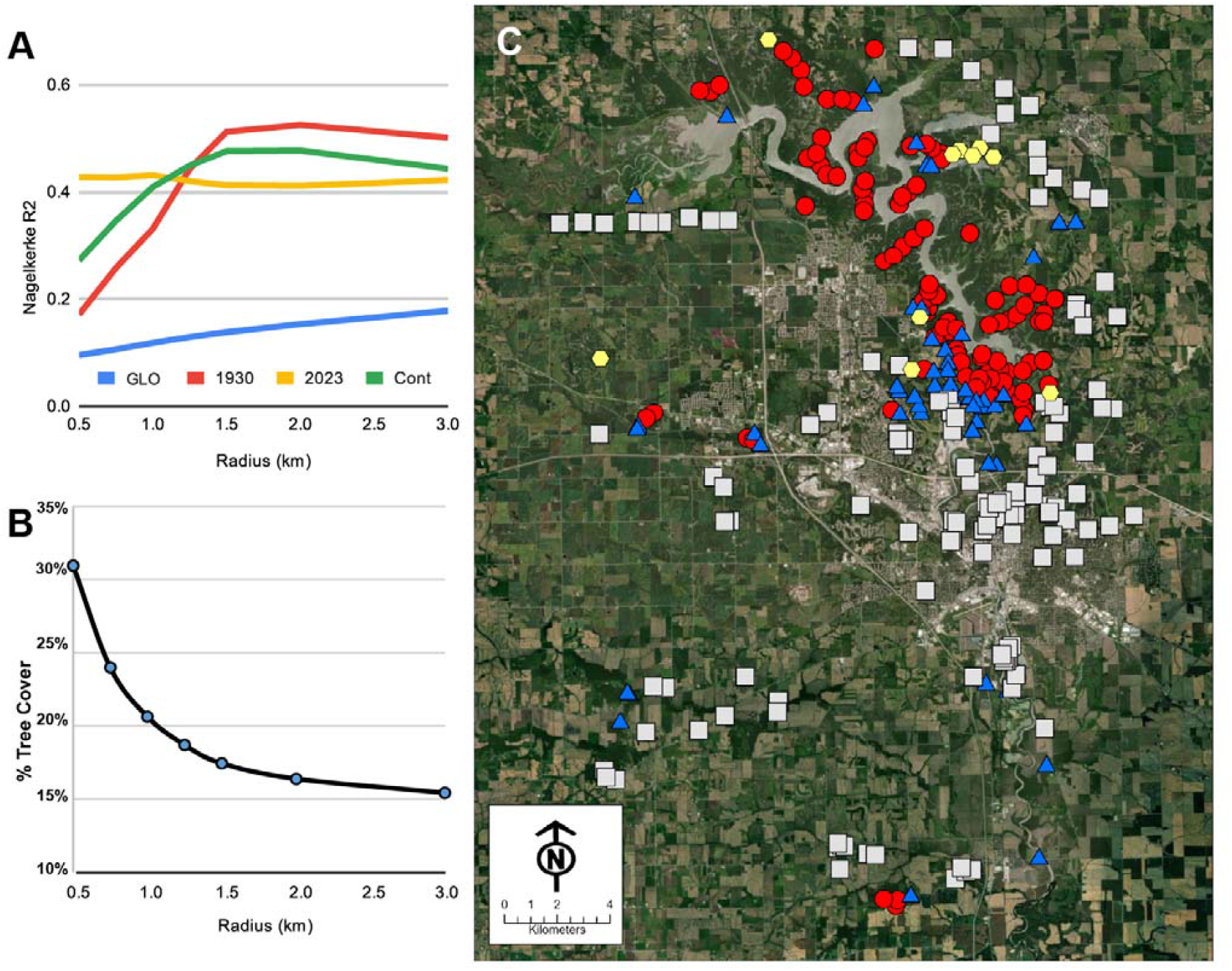
Legacy effects of forest loss and regrowth on *Magicicada* ranges. **(A)** Presence of chorusing cicadas at a site in 2024 was best predicted by the proportion of land in tree cover in a 1.5 – 2.0 km radius around the site in 1930. Tree cover in 2023 was the best predictor of cicadas only at very local scales (0 – 1 km). Forested habitat prior to considerable development (GLO; 1836 – 1859) was the poorest predictor of cicadas, though still significant. (B) Percentage of continuous tree cover (1930 – 2023) predicting a 90% chance of *Magicicada* presence at different radii. (C) Estimated change in cicada presence from 1930 – 2024 at 301 sites surveyed for chorusing in 2024. Presence of cicadas in 1930 was estimated using the 17.4% tree cover threshold of cicada suitability (Fig. 2C) applied to 1930s tree cover at 1.5 km. Red circles = cicadas predicted in 1930 and present in 2024; Blue triangles = cicadas predicted in 1930 but absent in 2024; White squares = cicadas not predicted in 1930 and absent in 2024; Yellow hexagons = cicadas not predicted in 1930 but present in 2024.

### Forests have expanded, but cicada ranges have not responded in kind

We used continuous tree cover, representing only those pixels of the map that had trees in both 1930 and 2023 (63.6 km^2^; 3.9% of the county; Figure S4) as an estimate of the amount of tree cover sufficient for long-term cicada persistence (Supplemental Methods). At a radius of 1.5 km (7.07 km^2^), binary logistic regression showed a 90% chance of cicada presence if ≥17.4% of the surrounding landscape had remained continuously in tree cover (Figure 2B). Applying this 17.4% threshold to the 2023 map suggested that 247 sites had sufficient tree cover to support cicada populations (Figure S5). This included 94.8% (110/116) of sites that had cicadas in 2024, with cicadas found at only six sites that the model suggested should not be able to support them. A further 137 sites were estimated to have sufficient forest cover to support cicadas and yet had none (Figure S5). These sites generally represented locations where trees had been removed prior to 1930 and then reestablished between 1930-2023 (Figures S2–S3).

Applying the same 17.4% tree cover threshold to the 1930s map offers an estimate of where *Magicicada* likely occurred in 1930, allowing comparison with 2024 chorusing data to visualize losses and gains between 1930-2024. Despite the increase in forest cover in the 94 years since 1930, more sites were estimated to have lost cicadas (55 sites) than gained them (10 sites) (Figure 2C). This recapitulates the same surprising retraction of cicada ranges in the face of a general increase of forested land that has been reported by several multi-generation *Magicicada* mapping projects (Cooley et al., 2004, 2013, 2016; Gilbert & Klass 2006).

The finding that cicadas are lost even as tree cover increases is explicable if one recognizes that increases in tree cover across a landscape are almost always net increases, inclusive of some turnover (Figure 3A). To wit: tree cover in Johnson County increased from 181.1 km^2^ to 206.7 km^2^ between 1930 and 2023, but only 63.6 km^2^ of tree cover was continuous from 1930 to 2023 (Figure S4). Thus, though the average net increase in forest cover was +0.27 km^2^/yr, this obscures an average tree cover loss of -1.01 km^2^/yr, or -17.2 km^2^ of potential habitat per *Magicicada* generation. Assuming that tree cover losses include areas both occupied and unoccupied by cicadas, this regular turnover results in a retraction of actual cicada ranges even as total potential cicada habitat increases every 17 years.

**Fig. 3.**
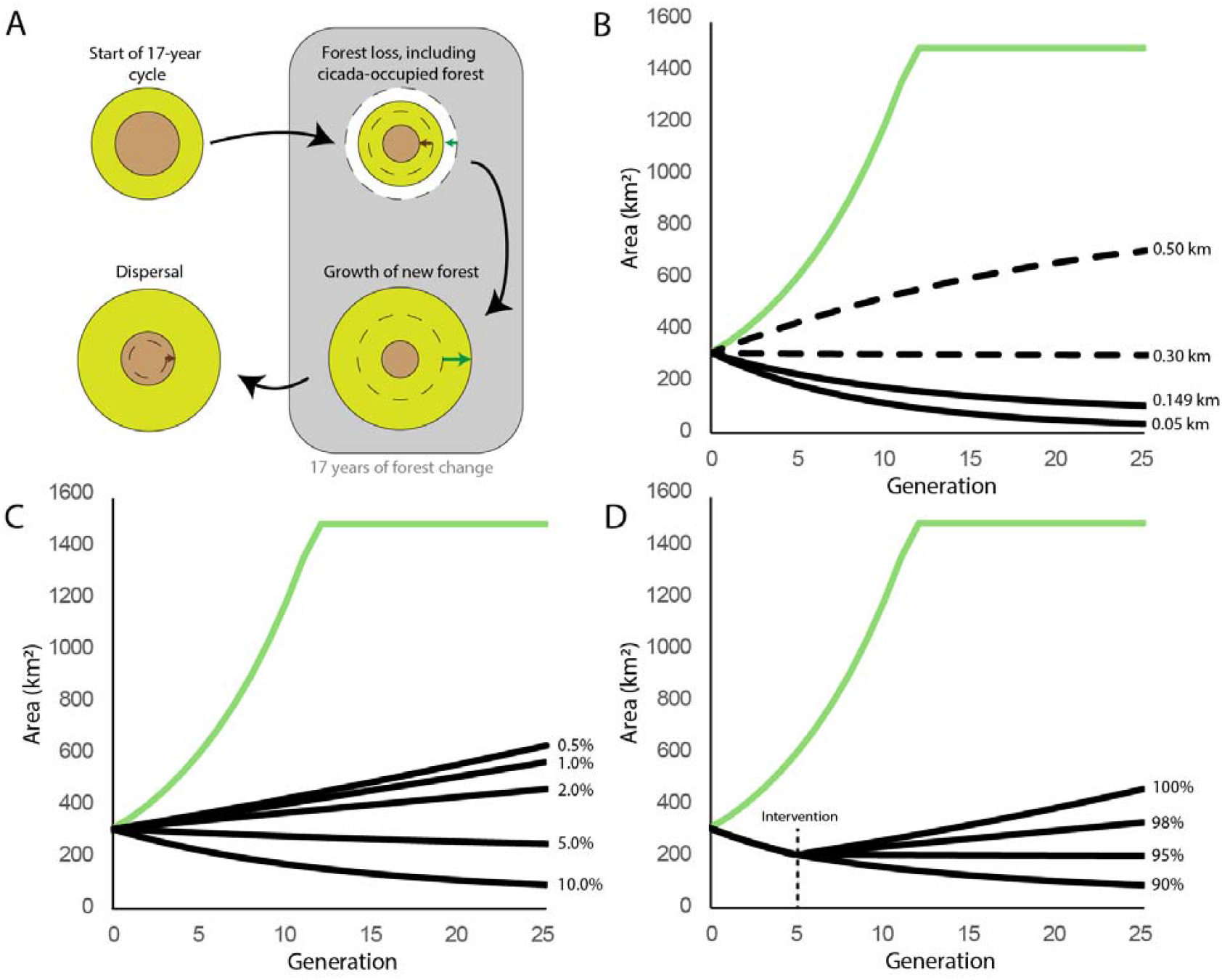
Why periodical cicada ranges decrease in a landscape even as tree cover increases. **(A)** The model imagines a contiguous circular area of forest occupied by cicadas (brown) within a larger circle of forest with no cicadas. During the 17 years cicadas are in the ground, some forest is lost resulting in the loss of cicadas and cicada habitat even if forest is added. At the end of each 17-year cycle, cicadas disperse into the surrounding forest. (B) Predicted changes in total tree cover (green) and cicada ranges across 25 generations (425 years) based on field observations of dispersal distances (solid black lines; Karban, 1981). Dashed lines = dispersal distances that would be necessary for cicada range persistence or growth. (C) Consequences of reducing forest turnover. If forest loss is kept below 4% per generation, cicada-occupied area could be maintained or slowly increase. (D) Consequences of intervening (after 5 generations) with targeted protections of cicada-occupied forest: If >95% of all cicada-occupied forest is protected from turnover, cicada ranges can increase. See Materials and Methods for details.

A model incorporating forest turnover and cicada dispersal demonstrates *Magicicada*’s sensitivity to forest change and explains our results (Supplemental Methods). With a loss of 11.9% forest and a gain of 14.5% new forest every 17 years (these percentages reflect the annual mean change across 93 years in Johnson Co.), total tree cover reaches a maximum in twelve generations (204 years; Figure 3B). Despite this, periodical cicada ranges continue to shrink under most realistic conditions. For instance, with a per-generation dispersal distance of 0.05 km (a generous lifetime travel distance for adult cicadas; Karban, 1981), cicada-occupied forest shrinks by more than half in seven generations (119 years). At the maximum measured dispersal distance for a mated female *Magicicada* (0.149 km; Karban, 1981), cicada-occupied forest is halved in 14 generations (Figure 3B). Even at a dispersal distance of 0.30 km, twice the recorded maximum, growth of cicada range is essentially flat, and only at unrealistic dispersal distances of 0.5 km and above do ranges grow (Figure 3B). If rates of forest turnover are set lower than the 11.8% recorded for Johnson County, cicadas fare better, but turnover must remain under ∼4% /17 yrs. to prevent ongoing cicada range reduction (Figure 3C). Similarly, a conservation effort focused on preserving cicada-occupied habitat could rescue cicadas, but only if >95% of all occupied habitat were to be indefinitely preserved without turnover, including new habitats into which cicadas disperse (Figure 3D). Even in these most favorable scenarios, expansion of *Magicicada* ranges by distances of just 1-2 km would take centuries.

### Relocations of cicadas to restored forests fail

Given the likely continuation of human-mediated forest turnover, *Magicicada*’s short dispersal distance is a critical impediment to their recolonization of restored forests (Figure 3B). To assess the feasibility of accelerating *Magicicada* dispersal, we conducted a managed relocation of adult cicadas to Pappy Dickens Preserve (“PD”). This managed forest in Iowa City had been reduced to pastureland sometime between 1860 and 1930 but had sufficient surrounding tree cover (40.8 % at a radius of 1.5 km) in 2024 to be characterized as an acceptable cicada habitat. PD was 3.8 km from the closest population of chorusing cicadas in 2024, and thus even if there were a fully forested corridor between sites with zero turnover, cicadas dispersing 0.05 – 0.149 km /generation would not recolonize PD naturally until sometime between the years 2466 and 3316. We relocated 20,126 adult cicadas to PD from other sites in Johnson County across the course of 10 days. Though relocated cicadas moved quickly into trees and undergrowth, bird and small mammal activity was heavy at the immediate site of cicada release and many predators were observed eating cicadas during and after each release. No adult cicadas were observed after the last release day, and the relocation resulted in no chorusing and no apparent egg-lay.

This failure to establish a new and sustainable *Magicicada* population echoes previous translocation attempts (Marlatt 1907, Alexander & Moore, 1962, Lloyd 1987) and is almost certainly due to predator aggregation. Birds are the primary predators of adult cicadas, with more than 80 species known to feed on them (Getman-Pickering et al., 2023). During an emergence, birds respond numerically, aggregating in sites of high-density emergence (Karban 1982, Leonard 1964), and change their behaviors to become more efficient at cicada handling (Steward et al., 1988). Diffusion of cicadas from the margins of existing ranges is possible because mobile predators (especially birds) in the immediate area become satiated, allowing *Magicicada* mating and range expansion via dispersal of mated females. But move even a large number of cicadas to a new site too distant from any existing population and the protection offered by having other cicadas nearby is lost as all nearby birds converge on that single site. Though the abundance of forest-adapted bird species may decrease as forested area is lost, bird species adapted to agricultural or urban landscapes also eat periodical cicadas (Getman-Pickering et al., 2023) so predator numbers may not decrease as forest habitat is reduced. This may also help explain why some sites with relatively old tree cover nevertheless show a pattern of shrinking *Magicicada* ranges (Maier 1982). If tree cover in the surrounding landscape is close to or below a lower threshold of 17.4%, or if forest habitat becomes too patchy, predator aggregation may also reduce the edges of existing ranges.

### Prospects and peril for periodical cicadas

A worrisome extension of this study is its relevance for the future of periodical cicadas, for which the specter of extinction has been raised (Simon et al., 2022, Cooley et al., 2002). Agricultural abandonment and accompanying forest regrowth in the Eastern U.S.A. (Ramankutty et al., 2010) may provide a panacea, but net increases in forested land still inevitably include turnover, and most forested areas within the range of *Magicicada* are <50 years old (Pan et al., 2011). Given that the only apparent prospect for recolonization after forest restoration is a 50– 150 m advancement of a “cicada front” every 17 years, forests need not only to be restored but protected, with little to no turnover, across multiple centuries (Figure 3C,D). It thus seems more reasonable to conclude that the large-scale removal of North American forested land more than a century ago may have set *Magicicada* on the same surprising decline towards inevitable extinction that befell other North American animals of extreme abundance.

## Acknowledgments

Collections and Surveys: J Alfieri, C Berggren, A Brummett, G Brummett, N Brummett, A Caraballo, N Cole, M Fallon, AR Forbes, S Forbes, O Giannakouros, M Giannakouros, S Giannakouros, J Grime, E Jalinsky, J Jalinsky, M Jalinsky, C Johnson, A Kitchen, C Kitchen, O Kitchen, M Land, S Lawinger, K Long, H Martinez, C Mulcahy, B Najev, M Neiman, B Parent, D Roberts, E Robinson, A Stallman, J Taylor, P Taylor, N Timmer, J Westin, M Wittmer, J Young, M Weinberger, S Zuhlke. Collection permissions and permits: Bur Oak Land Trust, Johnson County Conservation Board, Iowa Department of Natural Resources. GIS contributions: Bret Barschak. Historical Newspaper Search Assistance: C Bendixen. Translation of 1800s Czech-language newspaper articles: K Bubeníková. Additional helpful advice/comments: H Sander, S Secchi. Special thanks to the Bur Oak Land Trust for permission to relocate cicadas into one of their forest preserves, and to JR Cooley for sharing his thoughts and advice regarding the fruitlessness of such relocation efforts. This project was self-funded, but material and institutional support was provided by the University of Iowa. Stipends for IC, FC-S, and JSC were provided by an NSF REU site grant to AAF (DBI 2149361)

## Author contributions

The concept for this study was developed by AAF, with considerable contributions from IC, SDH, CLW, GEB, MJG, and MJS. All authors contributed to data collection and analysis. Data visualizations were led by AAF, IC, AMS, SKM, and CH. Writing was led by AAF and IC, with all authors contributing to reviewing and editing.

