## Supplemental Methods, Figures, and Tables for "Periodical cicadas suffer legacy effects of century-old forest removal"

Study Area and Historical Evidence of Cicadas

This study was conducted in Johnson County, a 1610 km^2^ area of land in the southeastern quadrant of the state of Iowa, USA and near the western edge of the range of Brood XIII periodical cicadas. Johnson County was established in 1837 (a Brood XIII emergence year; 11 cicada-generations prior to the 2024 emergence). In 1838, Iowa became a U.S. Territory, and the following year (1839) a plat map was drawn out for the intended capital of the territory, Iowa City, which at the time was mostly undeveloped land (Schwieder 1996). A natural resources and vegetation survey conducted by the U.S. General Land Office (GLO) between 1836 and 1859 shows most of the county as prairie with large contiguous areas of “timber,” “grove,” or “oak barrens”, the latter of which extended down into and through Iowa City (fig. S1).

The first aerial photography of the county (1930) shows a changed landscape: a patchwork of farmland and fields with relatively small areas of contiguous tree cover (fig. S2). Most remaining forest in the county at the time was second and third growth (Jenkinson 1969) and relegated to the northeastern quadrat of the county, close to the Iowa River. In the late 1930s, a tributary of the Iowa River was dammed, forming Lake MacBride (Jenkinson 1969), and in 1958, the Iowa River itself was dammed, forming Coralville Lake (Bunker & Witzke 1987). Both lakes displaced additional tree cover, but Lake MacBride became an Iowa State Park, and the U.S. Army Corps managed much of the forested land around Coralville Lake, thus protecting forests in these areas from considerable further development. Tree cover in other parts of the county slowly recovered, especially in suburban areas north of Iowa City. Tree cover in the county reached ~12.8% by 2023 (fig. S3). This increase is reflective of a state-wide trend of forest area increase post-1974 (Jungst et al. 1998).

Johnson County was specifically noted by Marlatt (1907) as being home to Brood XIII periodical cicadas, and they remain in the county today. A 2007 survey by Cooley et al. (2013) recovered multiple *Magicicada* chorusing sites in and around areas close to the Coralville Lake and MacBride State Park. No other peer-reviewed records of periodical cicada emergences at specific sites in Johnson County are available, but emergences that occur in and around human habit are often newsworthy. Thus, we reasoned that if historical cicada emergences had occurred in Johnson County, IA - and especially in its largest city, Iowa City - there would have been mention of them in local newspapers of record. We searched digitized records of newspapers published in the city of Iowa City, IA between 1856 and 2007 for the search terms “cicada”, “cicadas”, “Brood XIII” and “locust.” We conducted a broad search but focused especially on stories written during nine Brood XIII emergence years: 2007, 1990, 1973, 1956, 1939, 1922, 1905, 1888, and 1871. Articles from Brood XIII emergence years that were clearly referential to the Rocky Mountain locust, *Melanopus spretus* (which went extinct by 1902), were filtered out. We made notes about the focus of all *Magicicada* stories, including whether they were anticipatory or documentary, and whether they specified emergence location(s) (data: Chan et al. 2025).

Tree cover classification

Based on land cover descriptions from the historical General Land Office (GLO) vegetation survey, we reclassified the data into two categories. The “tree” classification included areas labeled as “Timber,” “Oak Barrens,” and “Grove.” All other classes were grouped into a single “no trees” category. We identified the historic boundary of Iowa City using digitized maps from the city’s founding. These maps were georeferenced using both modern and historical street networks and used to define a bounding polygon representing the city's historical extent (fig. S1).

1930 county imagery was obtained from Iowa State University Geographic Information Systems Support & Research Facility with a 1 m spatial resolution and a single spectral band. These aerial photos were acquired by the US Department of Agriculture from 1936 to 1941 through a project funded by the Iowa Department of Natural Resources and the USDA Natural Resources Conservation Service. The georeferencing of the 1930s imagery had an error of greater than 5 m and was corrected using over 500 tie points with 2003 imagery from the USDA’s National Agricultural Imagery Program (NAIP). The mean and variance across a 3x3 m moving window were calculated using the single band. The spectral band, mean, and variance product were used for 1930 tree classification. The 2023 imagery was obtained from NAIP and had a 0.6 m spatial resolution with four spectral bands (e.g., red, green, blue, and near infrared). The NAIP tiles comprising Johnson County were resampled to 1 m spatial resolution to match the 1930 spatial resolution and mosaicked together. The variance across a 3x3 m moving window was calculated using the near-infrared band. After determining which bands contained the most spectral discrimination, the red band, near infrared band, and variance product were used for 2023 tree classification.

Training sites were developed by distributing 500 random points within the Johnson County boundary. Using the aerial imagery as a background, each site was labeled either ‘tree’ or ‘other’ for both 1930 and 2023 imagery. For each site, the closest pixel with the smallest Euclidean distance from the site was extracted from imagery. We employed a binary support vector machine (SVM) classification to classify pixels containing trees for each year. Using all sites, the SVM binary classification parameters were optimized using a Bayesian optimization approach with 300 objective function evaluations, which determined the optimal box constraint, kernel function, kernel scale, polynomial order (if the kernel function is polynomial), and whether data standardization improves results. After determining the optimal parameters, SVM models were trained over 100 iterations, with each iteration randomly selecting 70% of sites within each class (tree versus absent). The confusion matrix is calculated using the remaining 30%, and the average overall accuracy, producer accuracy, user accuracy, and kappa are reported. To develop the final image classification, 25 SVM models (randomly selected) were run on the image. Each pixel’s mode from the 25 iterations determines the final classification value. Classification confidence is calculated as the percentage of times a pixel was classified as the final classification class. The final tree classification used for analysis was determined by selecting all pixels classified as trees with a confidence of over 90%. The confusion matrix was calculated using this final classification image and average overall accuracy, producer accuracy, user accuracy, and kappa are reported (table S2). Final maps are shown in figs. S2-S3.

The above process resulted in two raster datasets for 1930 and 2023, a classification with tree presence/absence (0/1) and a classification confidence with a "confidence" rating (0 – 25). Using raster calculator, we selected pixels that were classified as tree (1) and scored the highest (25) on the confidence scale. To determine continuous cover, we used the raster calculator to find pixels that were classified as tree with high confidence in both 1930 and 2023. To remove water from the 2023 and continuous datasets we used the water bodies layer from the USGS National Hydrography Dataset (NHD). Maps are shown in figs. S2-S4.

We constructed buffers around each of the 301 sites of 0.5 km, 0.75 km, 1 km, 1.25 km, 1.5 km, 2 km, and 3 km. We then ran a zonal statistics analysis to count the trees within each buffered area. Since cells were 1 m x 1 m, the number of cells is equivalent to the area of tree cover (in m^2^) for each buffered area. This was done for all three datasets, 1930s, 2023, and continuous.

Survey of 16 forested areas for *Magicicada*

Between April 25th and May 10^th^, 2024, prior to the cicada emergence, we surveyed 16 forested sites across our study area (Fig. 1; Table 1) for pre-emergence holes. Each hole represents one cicada (Dybas & Davis 1962), and thus hole density can be used as an approximation of cicada density. Fourteen sites were within the area identified as having had tree cover in the 1836-1859 GLO map (Fig. S1), with the other two sites (sites #6 and #15) outside this area. At each site we haphazardly selected between two and 37 trees (mean = 6.6; median = 5) sufficiently large that we could be sure they had been present for at least one and likely several *Magicicada* generations. We recorded the GPS coordinates of each tree using the GAIA GPS app (Outside Interactive, Inc., Boulder, CO) and counted the number of emergence holes in five 1 m x 1 m quadrats (fig. S6A) placed around the tree ~1–2 m away from the trunk. To expose cicada emergence tunnels (fig. S6B), we removed any leaf litter by hand and scraped away the top ~3–5 cm of soil with the long edge of a hand trowel. We counted all holes larger than 1.0 cm to exclude earthworm tunneling, though we acknowledge that some holes that we counted could represent activity of ground-nesting bees or other animals. In some holes a cicada nymph was visible (fig. S6C). We averaged the number of exit holes / tree across the five quadrats sampled, and then again across all trees for a mean of means (Table 1). For most, but not all, trees we also made notes about the tree species, notable characteristics, and surroundings, including vegetation. Raw data: Chan et al. (2025).

We returned to each of the 16 sites in June during and after cicada emergence to look for evidence of an emergence. Adult periodical cicadas emerge in large numbers across consecutive nights (Williams & Simon 1996) and newly-emerged adults remain on low-lying vegetation for much of each morning (figure S6D), so presence or absence of a cicada population is easy to assess, requiring simply a visit to the site. We also monitored the crowd-sourced apps iNaturalist and CicadaSafari during this time to see where other people observed cicada emergences at or near any of the same 16 sites. Cicada mass emergences were only observed at sites #1, #2, #3, #4, and #5 (Table 1). At one additional site (#9) we observed five shed *Magicicada* skins but did not see or hear adults. We saw no additional evidence of adult cicadas when visiting any of the remaining ten sites, but contributions by users of naturalist phone apps showed evidence of extremely-low density cicadas at three other sites: a shed skin (posted by iNaturalist user *Konung-yaropolk* on May 27) and a single adult (*mafri* on May 31) at site #9, an adult *M. cassini* within a few meters of site 13 (*rhinogradentia* on June 3), and an adult at site 16 (a user of the CicadaSafari app).

We also revisited the same sixteen sites to listen for male chorusing and assess evidence of damage to young tree branches caused by cicada oviposition. Just as an established population of cicadas is easy to visually assess as they emerge, chorusing of male periodical cicadas is loud (reaching 80 dB; *34*) and is trivial to detect without additional equipment (Video S1). Males require at least 5 days after emergence to begin chorusing (Williams & Simon 1996), and during this time pressure from bird and mammal predators is extremely high (Karban 1982; Williams et al. 1993). Thus, it is likely that chorusing only occurs when cicada population sizes are sufficiently large to have satiated local predators. Chorusing therefore presents a clear and obvious signal of a well-established local population and is one that has been used in previous mapping work (e.g., Cooley et al. 2021). After mating, heavy oviposition by *Magicicada* females causes tips of branches to wither, a visually obvious phenomenon termed “flagging.” We only heard sustained chorusing and observed “flagging” of branches at sites #1 – #5, though a team member heard a single male *M. cassini* call at site #10 that was not repeated.

*Magicicada* chorusing surveys

On sunny days between mid-morning and early afternoon (mostly 9 am to 2 pm) during peak chorusing weeks (May 27 to June 20), we drove to forested sites across the study area and listened for nearby chorusing. We categorized each site as either having chorusing cicadas or not. The three *Magicicada* species emerging in Johnson Co., IA (*M. cassini*, *M septendecim*, and *M. septendecula*) have different male mating calls that can be easily distinguished from one another. We did not uniformly spend time distinguishing among calls of the three species but often noticed at least two distinct call types (usually *M. cassini* and *M. septendecim*) at all “loud” sites. We plotted each point on the Gaia GPS app (Trailbehind, Berkeley, CA) in real time while surveying and later transferred the data to an ArcGIS Pro project. We recorded chorusing at 116 of the 301 locations surveyed (data: Chan et al. 2025).

Models

We ran binomial logistic regressions in R Studio to assess the relationship between tree cover density (1860, 1930, 2023, continuously-forested) and cicada presence in 2023. We tested seven different buffer radii around each point (0.5, 0.75, 1.0, 1.25, 1.5, 2.0, and 3.0 km; data: Chan et al. 2025). To account for large bodies of water (Lake MacBride and Coralville Lake) that would otherwise reduce the total land cover, we computed the fraction of tree cover in 2023, and continuous tree cover as (tree cover area) / (total area in radius - water area in radius). For 1930 and GLO forest cover, we did not subtract the area of water from the total area as those lakes did not exist at those times. We used the binomial variable of cicada presence in 2023 at the 301 survey sites as the response variable and the tree cover densities as the explanatory variables. We used the function lrm from the rms package to calculate the Nagelkerke R^2^ value and glm function from the base stats package to calculate the AIC value. We used these values to assess the predictive power and level of fit at each radius. We also combined multiple tree cover densities at different time points and radii to assess how these affected the model. Results are found in table S3.

To determine the proportion of tree cover necessary to support cicadas across generations, we determined the level of continuous tree cover (1930-2023) that resulted in the highest likelihood of cicadas in 2024. With the logic that this level of continuous forest cover represents the tree cover required to sustain cicada populations across long periods of time, we used these results to predict which areas of Johnson County in 2023 had sufficient forest cover to support cicada populations (regardless of 2023 cicada presence). After obtaining variable coefficients and intercepts, we plotted logistic graphs and found the x-values (tree cover density) at which the y-values (probability of cicada presence) equaled 0.9 to represent 90% confidence. We then determined which areas in 2023 had forest cover greater than these cutoff values to predict which forests were likely able to support cicada populations (fig. S5).

To estimate where periodical cicadas may have been in 1930 and subsequent loss or gain of territory, we used the same cutoff for habitat suitability based on continuous forest density at 1.5 km at 90% confidence. We then compared sites predicted by the model to have had cicadas in 1930 to chorusing data from 2023 (Fig. 2C). Estimating that more sites had lost cicadas (n = 55) than gained them (n = 10) since 1930, despite a ~14% increase in tree cover during that same 93-year interval, we made a model of habitat change to explain how forests can expand while cicadas ranges decrease. To parameterize the model, we calculated average tree cover gained per year by taking the difference between 1930 and 2023 tree cover levels (+25.5 km^2^) and dividing by 93 to get an annual net change of +0.27 km^2^. Because this represents a net gain and does not inform about how much current forest is lost per generation, we also calculated an annual rate of turnover by taking the difference between 1930 forest cover and continuous forest cover (-117.6 km^2^) and again dividing by 93 = -1.26 km^2^. Finally, we added 1.26 to 0.27 to estimate gross annual forest gain (1.54 km^2^ / yr). We multiplied annual turnover rate and gross tree cover gain both by 17 generations and divided by 1930 tree cover to estimate the proportion of tree cover lost (11.9%) and gained (14.5%), respectively, in a 17-year cicada generation. We do not assert that these are precise estimates, and for that reason tested several additional (and more conservative) variations (see below).

Our model (Fig. 3A; Chan et al. 2025) imagines a contiguous, circular forested area with a radius of 10 km (314.16 km^2^) in a larger landscape of 1500 km^2^. We used different areas to track the size of total forest and cicada-occupied forest, but at the start of generation 0, cicadas occupy all forested area, so the two areas are initially the same. Both areas then lose 11.9% of their area followed by a gain of 14.5% of the non-cicada forest area, representing forest change across a 17-year period. Importantly, the cicada-occupied area does not increase during this period because cicadas must disperse in order to occupy new forest. The area of cicada-occupied forest increases in the model by the addition of a cicada dispersal distance to its radius. Cicada-occupied forested area cannot exceed total forested area, and total forested area cannot exceed 1500 km^2^. Subsequent generations use the same sequence of events: subtraction of 11.9% from both areas, addition of 14.5% to total forest, and finally cicada dispersal to increase the area of cicada-occupied forest (Data S1).

We tested four dispersal distances (0.05 km, 0.149 km, 0.3 km, and 0.5 km). Cicadas travel extremely short distances across their adult lifespans, mostly staying in or near the trees where chorusing occurs (Marlatt 1907; Craig 1941; Karban 1981). Karban (1981) provides the most direct assessment of dispersal distances, using mark-recapture and direct observation to find that 90% of *Magicicada* adults moved <50 m across their lifespan, <1% of adults flew more than 100m, and a single mated female flew 149 m. Cooley et al. (2021) indirectly inferred dispersal distances potentially as large as 1-2 km, but the authors acknowledged that the spatial resolution of sampling was such that these movements could have been considerably shorter. Inclusion of 0.3 km and 0.5 km distances test the consequences of longer dispersal distances than those that have been directly measured in nature.

Because loss of cicada habitat during each 17-year period has a major impact on range reductions, we tested how different percentages of forest turnover affected outcomes. We adjusted the Johnson County rate of 11.9% / 17-years to 10%, 5%, 2%, 1%, and 0.5%. We separately simulated a scenario whereby after five years of decline, an intervention resulted in protection of all or most cicada-occupied habitat (including newly colonized habitat). We tested the consequences of protecting 100%, 98%, 95%, and 90% of cicada-occupied forest. For all forest turnover variations we used a dispersal distance of 100 m, splitting the difference between 50 m and 149 m. We also simulated outcomes with a 13-year cicada life cycle and with an annual cycle. Results of model outcomes: Fig. 3B-D, Data S1.

Managed relocation of adult cicadas

Discovering that cicadas have declined even as tree cover has grown in Johnson County, and that the prospects for natural recolonization were poor (see main text), we assessed the feasibility of reintroducing cicadas to land they had previously inhabited and that our model suggested had acceptable tree cover to support cicadas. We identified a restored forest site within the area of GLO land classified as having trees in the 1860s and that had lost cicadas after being reduced to pastureland between 1850 and 1930, but had the characteristics, in 2024, of an acceptable cicada habitat. The restored site, Pappy Dickens Preserve (part of site #9 in the preemergence surveys), abuts another mostly-forested site, Hickory Hill park. A circle of land with radius 1.5 km around Pappy Dickens and Hickory Hill contains 40.8% tree cover, such that the area in general fits the profile of “acceptable” cicada habitat (see Main Text).

Periodical cicadas are not likely to return to Pappy Dickens Preserve on their own in the near future. Assuming absolutely no forest turnover and a dispersal rate of 50–149 m / 17 years, a natural diffusion from the closest site where we found cicadas at high density (~3.78 km) would not return *Magicicada* to Pappy Dickens until 2466–3316. Thus, if a reintroduction were to be successful it would provide a blueprint for more rapid periodical cicada recolonization of conservation without the necessity of re-foresting or creating corridors between acceptable sites.

Transplants of periodical cicadas have been attempted before and have not previously resulted in reestablishment of breeding populations. A transplant of ~1000 adult cicadas was entirely consumed by birds in three days (Alexander & Moore 1962). However, we found no evidence that more than 1000 adult cicadas had previously been moved (a reference to 10,000 adults moved by Martlatt [Karban 1982] appears to be in error). Two other efforts employed the movement of egg-laden branches that introduced an estimated tens of thousands to hundreds of thousands of nymphs into soils (Marlatt 1907, Lloyd 1987). Both cases also failed, with many nymphs being consumed by ants and all resulting adults in the next generation being eaten by predators. Though the evidence seemed weighted against such efforts, predator satiation appears to occur when cicada densities are ~24,000/ha (Williams et al. 1993), and thus we wondered whether an extremely concentrated introduction of 20,000 cicadas might overwhelm local predators.

We requested and received *Magicicada* collection permits and permissions from the Iowa Department of Natural Resources (permit SC1553), the Johnson County Conservation Department (permit 042624), the US Army Corps of Engineers, and the Bur Oak Land Trust. We collected cicadas at five sites in Johnson County, IA: Big Grove Preserve, Cangleska Wakan, Two Horse Farm, Turkey Creek Preserve, and MacBride Nature Recreation Area (MNRA) (sites 1-4 in Fig. 1). All five sites had pre-emergence holes in early May 2024, and what we estimated to be extremely high *Magicicada* densities (this estimation was later proved out by observations of several weeks of extended adult chorusing, mating, and egg-laying; Table 1)

We also received permission from the Bur Oak Land Trust to relocate cicadas to Pappy Dickens Preserve. This was not a particularly long-distance relocation: Pappy Dickens is just 7.4 km from the nearest source site (Turkey Creek Preserve) and 14.0 km from the farthest source site (MNRA). Pre-emergence cicada hole surveys at Pappy Dickens found no holes, and the same surveys at the adjacent Hickory Hill Park forested areas suggested extremely low densities of cicadas (just three holes across a total of 31 sample locations). We later observed only five shed cicada exoskeletons during daily walks of Pappy Dickens and none at Hickory Hill, and saw no adult cicadas (though we noted an iNaturalist observation of a single adult periodical cicada at Hickory Hill on May 31st). In short, we observed no sign of a mass periodical cicada emergence from soils in either area.

We fashioned sixteen collection containers by cutting a 10.5 cm diameter hole in the lid of a plastic tub (51.5 cm x 34 cm x 13 cm) and cutting the top off of a small deli cup. We then glued the deli cup into the hole. The lid of the deli cup could be removed to add cicadas but its protuberance below the container lid effectively prevented cicadas from flying or walking out. We attached collection instructions to the lid, including pictures to help collectors identify and exclude *Massospora cicadina*-infected cicadas.

Collections were made from May 24th, 2024 to June 2nd, 2024 by the authors as well as >20 local volunteers. We included volunteers on an email list that shared the goals, methods, and possible outcomes of the project. Collections were made primarily in the morning, between the hours of 8am and noon, when cicadas had only recently shed their nymphal exoskeletons and most had not yet made their way into tree canopies where they would be inaccessible. Collection containers were never filled much beyond 1000 adults; even at those largest numbers we observed only 1-2 deaths per container between their collection and release later the same day.

Collection containers full of periodical cicadas were transported to the Hendrix lab at the University of Iowa, where approximately 1/10th of each collection was subsampled to determine relative ratios of each of the three *Magicicada* species, as well as sex ratios of each. Five collections totaling 729 cicadas of the 20,126 live cicadas released were not subsampled for sex ratios or species IDs. We released all cicadas in a 20 m x 20 m area of Pappy Dickens, reasoning that this would be the best way to achieve large local densities. Approximately half of all released cicadas were deposited by hand on the lower trunk or base of one of four trees in the release area, while the remaining half were either scattered in low-lying vegetation or flung into the air (the latter strategy inevitably led to the cicada flying to a nearby tree or shrub).

For the first six days, cicadas were captured and then released at Pappy Dickens on the same day as their collection. By May 30, with no sign that any cicadas had persisted at the release site, we decided to switch to releasing older cicadas that would be closer to reproductive maturity. Thus, cicadas collected on May 30 and 31 were held in an insect cage constructed from mesh and duct tape in a walk-in incubator. We provided these cicadas with potted hardwood tree saplings (hackberry, red oak, walnut) and cut branches in water. Though individuals that fed on branches and trees survived well in the lab, many of the cicadas that crawled on the sides and ceiling of the enclosure died. On June 2nd, we released the surviving 4,677 adults, and these plus an additional 70 collected that morning were the last adults to be released. We also added 5,078 dead cicadas to the release site, reasoning that they might help satiate ground-foraging animals.

In total, we released 20,126 live periodical cicadas into Pappy Dickens Preserve. Of the 2613 *Magicicada* that were subsampled, 1736 (66.4%) were *M. cassini*, 753 (28.8%) were *M. septendecim*, and 124 (4.7%) were *M. septendecula*. Across all species, the M:F ratio was 51.8% - 48.2%. Cicadas were generally active upon release, walking up trees and readily flying across or away from the release site (Videos S2, S3). Birds and chipmunks were also observed actively eating cicadas immediately following each release.

After each release event, one or more members of the research team stayed in Pappy Dickens for 0.5-1 hour to observe the activity of released cicadas, as well as the activity of birds and rodents. Every 1-2 days after releases had begun, one or more of us walked a path of trails around the release site (and including the release site) to listen and look for signs of cicada activity. As an *ad hoc* measure of bird predation (often the primary cause of cicada mortality; Karban 1982; Williams et al. 1993), on May 28 we marked three 1 m^2^ quadrats at the release site with flags, cleared them of leaves and vegetation, and counted (and then removed) all cicada wings. These quadrats were 0 m, 1 m, and 4 m away, respectively, from one of the four trees in the release site. We counted wings in each quadrat again on May 29^th^ and May 30th. Wing counts in the three 1 m^2^ quadrats offered a signal of consistent bird predation across all three days of data (data: Chan et al. 2025). We calculated the number of dead cicadas implied by our wing counts across the entire 400 m^2^ plot by taking an average number of wings per quadrat per day, dividing by four (cicadas have four wings), multiplying by 400, and adding the result from the three days together. This offered an estimate of 15,432 dead cicadas, just 17 cicadas short of the 15,449 that had been released by that point in the study. Though this estimate is imprecise, it demonstrates the general magnitude of predation at the site.

We continued to monitor the site daily for any cicada sign until June 12th (ten days after the final release) and irregularly for an additional month. Adult cicadas were not observed in large numbers the day after any release, with a maximum of three adults observed on low-lying undergrowth the day following a release. No chorusing or even single male cicada calls were heard at the site from May 24 (the first release day) to June 12 (ten days after the final release), except during the 1-2 hours immediately following a new release. Many of the 5,078 already-dead cicadas were also observed to be consumed in a 5-day period (data: Chan et al. 2025).

Jenkinson, Golda L. 1969. *A history of Lake MacBride State Park*. Solon, IA:Cottage Reserve Corp.

Jungst, Stephen E., Donald R. Farrar, and Michael Brandrup. 1998. “Iowa's changing forest resources.” *Journal of the Iowa Academy of Science* 105: 61–66.

Karban, Richard. 1981. “Flight and dispersal of periodical cicadas.” *Oecologia* 49: 385–390.

Karban, Richard. 1982. “Increased reproductive success at high densities and predator satiation for periodical cicadas.” *Ecology* 63: 321–328.

Lloyd, Monte. 1987. “A successful rearing of 13-year periodical cicadas beyond their present range and beyond that of 17-year cicadas.” *American Midland Naturalist* 117: 362–368.

Marlatt, Charles L. “The Periodical Cicada.” *United States Department of Agriculture, Bureau of Entomology Bulletin* 71: 1–183.

Schwieder, Dorothy. 1996. *Iowa: the middle land*. Iowa City: University of Iowa Press.

Williams, Kathy S. and Chris Simon. 1995. “The ecology, behavior, and evolution of periodical cicadas.” *Annual review of entomology* 40: 269–295.

Williams, Kathy S. and Kimberly G. Smith. 1991. “Dynamics of periodical cicada chorus centers (Homoptera: Cicadidae: Magicicada).” Journal of Insect Behavior 4: 275–291.

Williams, Kathy S., Kimberly G. Smith, and Frederick M. Stephen. 1993. “Emergence of 13‐yr periodical cicadas (Cicadidae: Magicicada): Phenology, mortality, and predator satiation.” *Ecology* 74: 1143–1152.

Fig. S1.

**Map of Johnson County, IA showing forested landcover (green) as recorded by the United States General Land Office (GLO) between 1836 and 1859**. The dashed lines indicate the original boundaries of Iowa City in 1839, and yellow dots indicate two specific locations of *Magicicada* emergences reported in Iowa City newspapers in 1888 and 1905, an indication that cicadas were present in these historical forests:

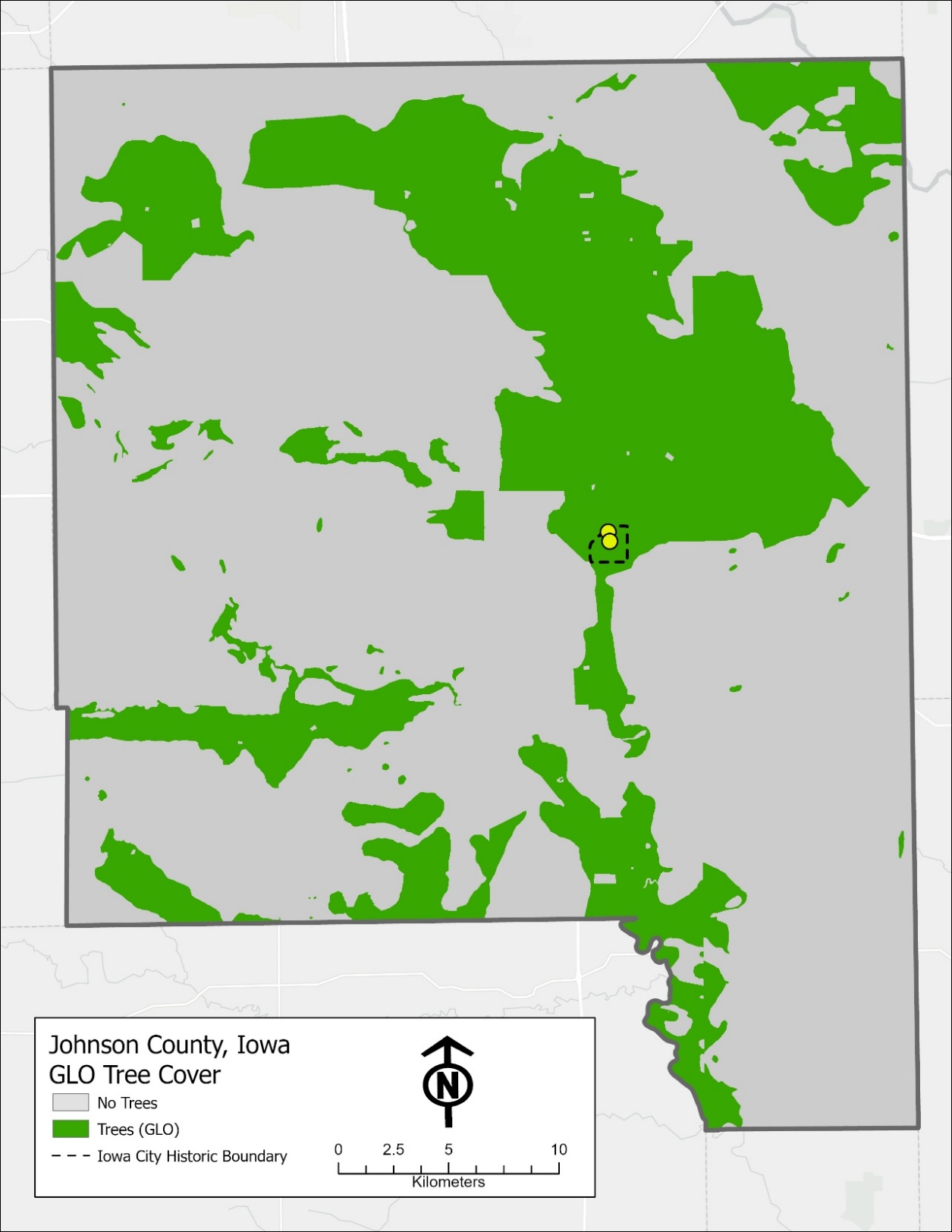

Fig. S2.

**Tree cover in Johnson County, IA in 1930.** White pixels indicate presence of tree cover at a 1m x 1m resolution.

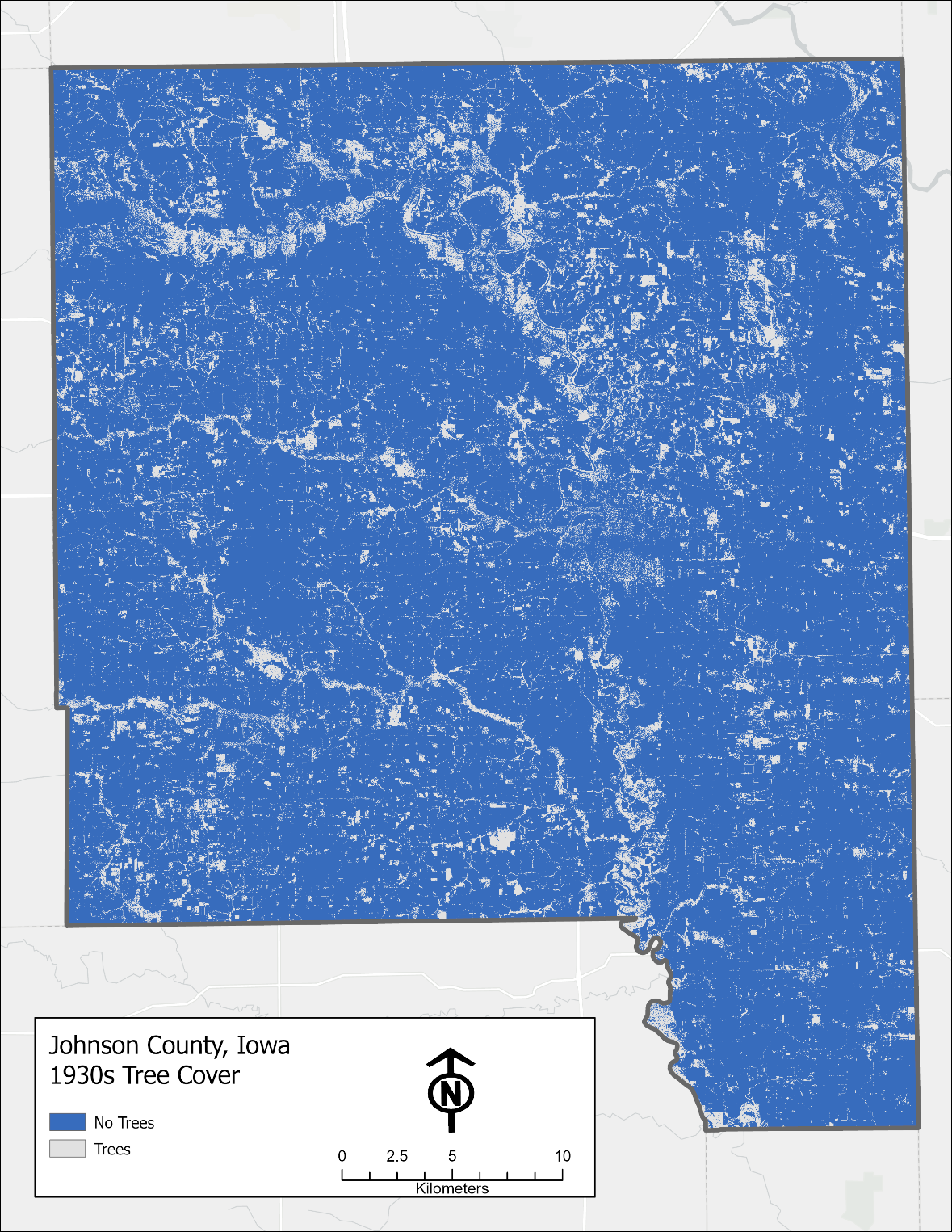

Fig. S3.

**Tree cover in Johnson County, IA in 2023.** White pixels indicate presence of tree cover at a 1m x 1m resolution.

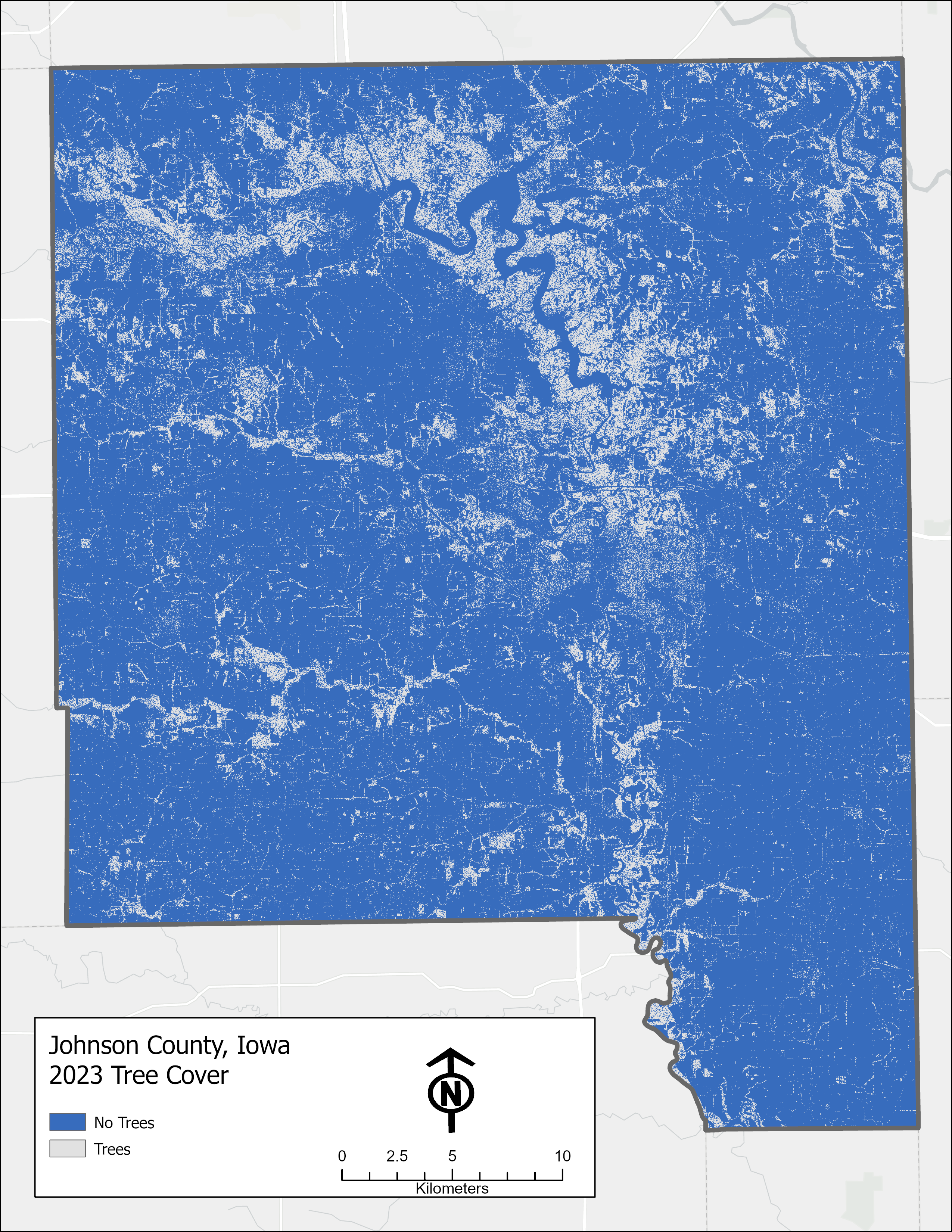

Fig. S4.

**Continuous tree cover in Johnson County, IA 1930-2023.** White pixels indicate presence of tree cover at a 1m x 1m resolution.
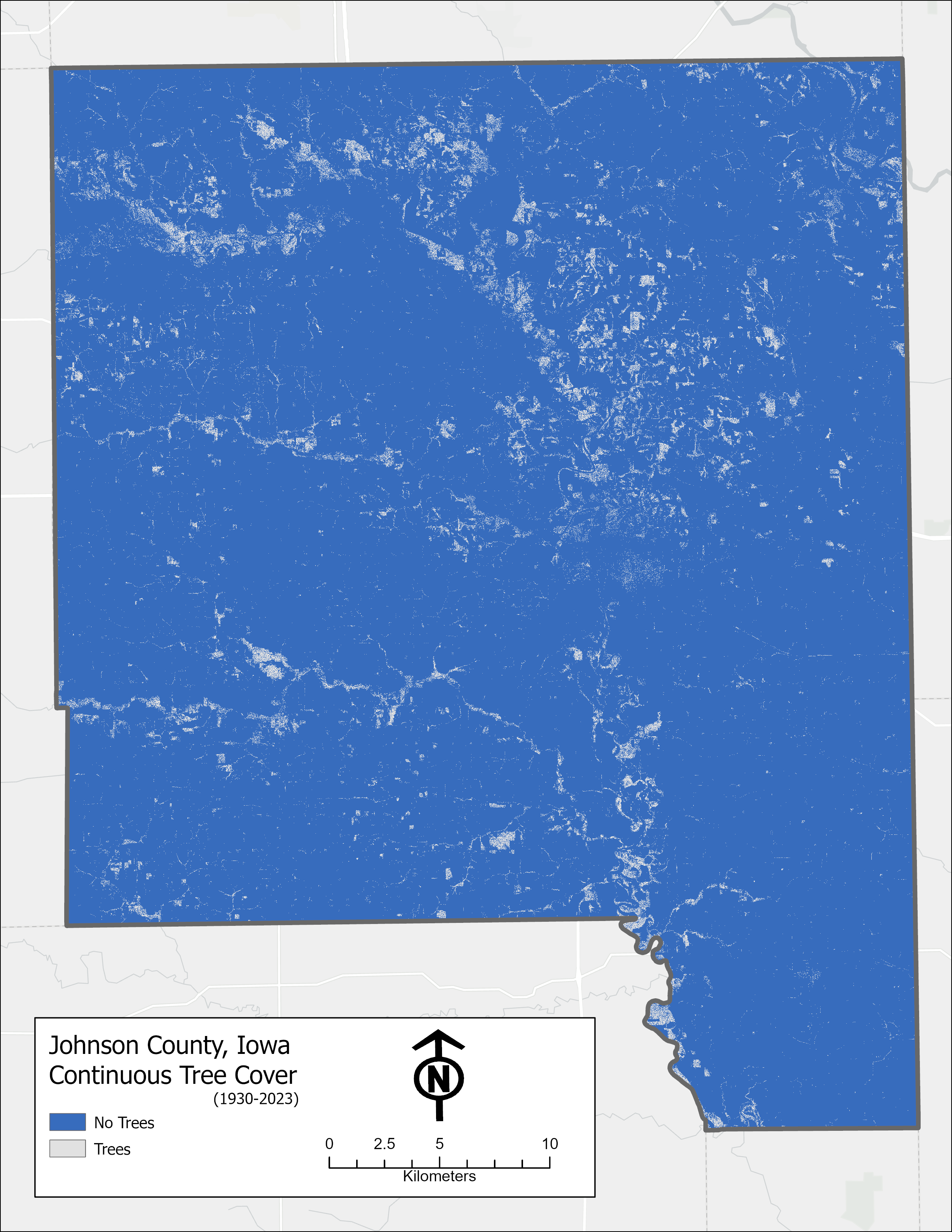

**Fig S5.**

**Sample sites with sufficient forest cover to be acceptable *Magicicada* habitat in 2023 versus where *Magicicada* occurred in 2024**. Red circles = acceptable habitat and cicadas present; Blue triangles = acceptable habitat but cicadas absent; yellow hexagons = not acceptable habitat, cicadas present; white squares = not acceptable habitat, cicadas absent.

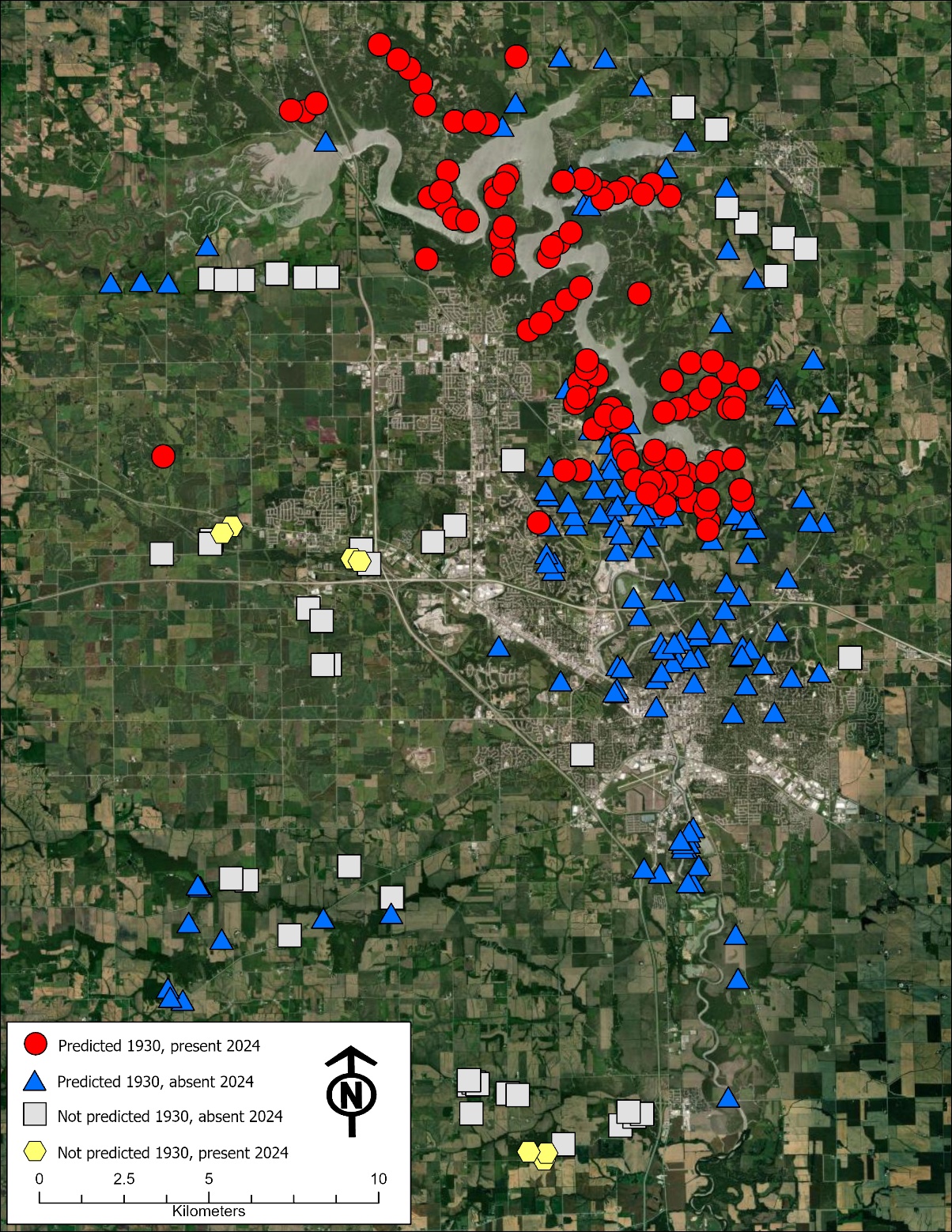

Fig. S6.

**Methods for documenting *Magicicada* presence at sites**. **(A)** 1 m x 1 m quadrat used to survey for pre-emergence cicada holes. (B) Close up of cicada holes with arrows indicating locations. (C) Cicada nymphs were visible in some holes. (D) On days that cicadas emerge from soil, adult cicadas and their shed cases are found on low lying vegetation across a site.
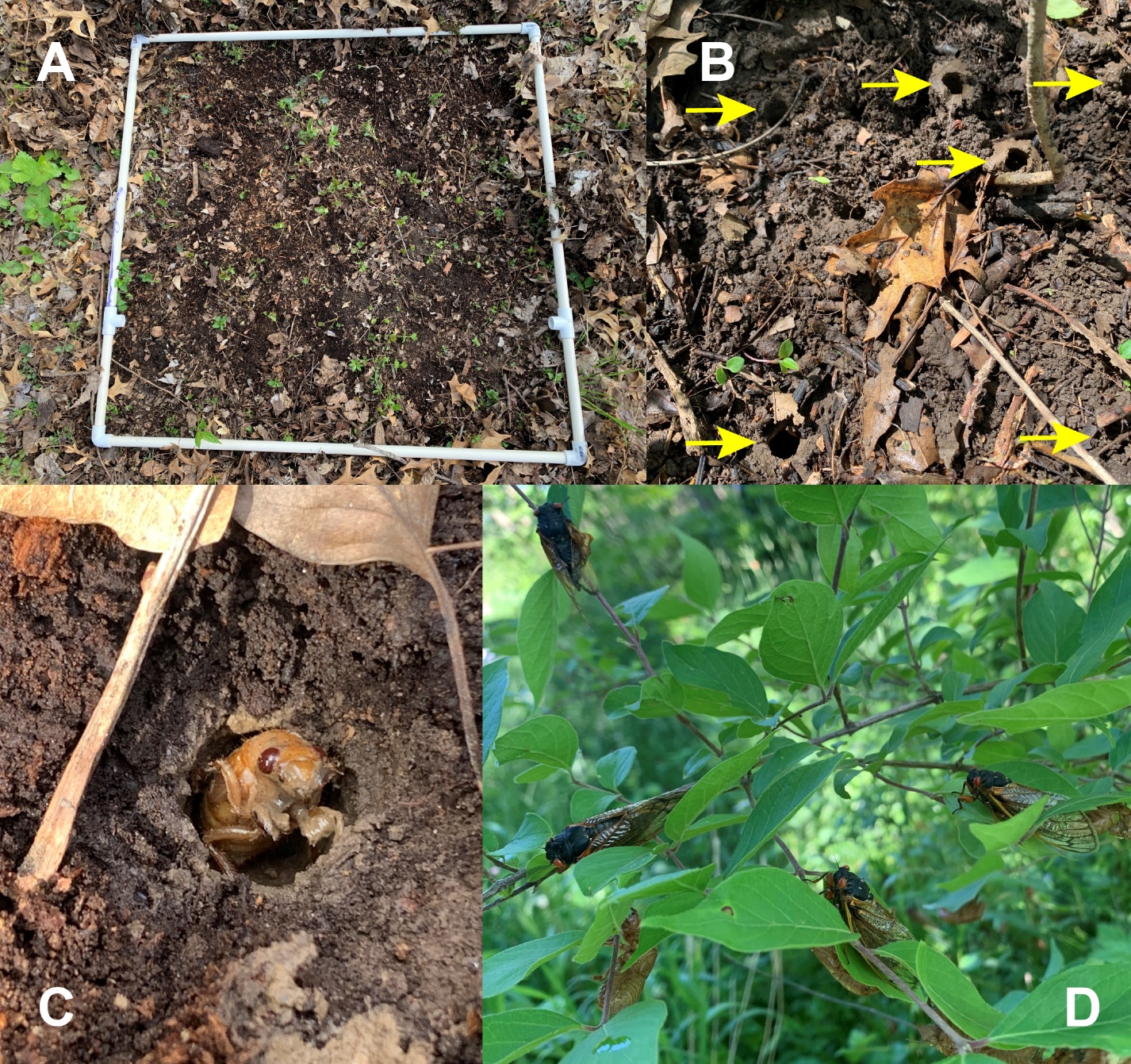

Table S1.

**References to Brood XIII periodical cicada emergences in Iowa City, IA.** Specific mentions of emergences in Iowa City are bolded and underlined. Though cicadas are referenced in future years, emergences in Iowa City are only noted in 1871, 1888, and 1905, after which references to cicadas are only anticipatory, or emergences are only described as occurring at sites elsewhere in Johnson County. Full data, including anticipatory mentions in later years, may be found on Dryad: <https://doi.org/10.5061/dryad.4tmpg4fq8>.

| Year | Newspaper | Mention of periodical cicadas |
| --- | --- | --- |
| 1871 | Iowa City Republican (1856-1921) | 1) "The Seventeen Year 'Locust'" June 18, 1871: A general information article about the habits of the periodical cicada. 2) Untitled article, June 21, 1871: uses the term "locust" but is clearly describing a periodical cicada emergence, as the Rocky Mountain locust would not have been laying eggs into fruit trees. Also has a clear reference to cicadas being in the city of Iowa City itself. "The locusts are now committing their ravages in the city, by puncturing the limbs of fruit trees and depositing their eggs there-in...It is almost a hopeless task to attempt to fight so many, but it is supposed they will not stay much longer and the trees are worth the saving." |
|  | Daily Evening Press (1871-1871) | May 6, 1871: this anticipatory note: "Seventeen year locusts are to make us a call this year." 2) June 8, 1871: this description of an emergence in the city: "The song of the locust is getting louder. They are invading the city" |
|  | Slovan Amerikansky (1870-1873) | June 15, 1871 (in Czech): Describes how, after a heavy rain that damaged roofs in Iowa City, periodical cicadas were present all around, and making lots of noise. The article also describes the morphology and life history of the 17-year cicada and distinguishes them from grasshoppers (locusts). |
| 1888 | Vidette-Reporter/Daily Iowan (1868 - 2024) | “Some observations on the Cicada”, by Minnie Howe, June 9, 1888: describes a mass emergence of cicadas "in a yard on North Dubuque Street" in Iowa City: "Trees, fence, grass and shrubs were covered with the cast-off pupa skins or the winged imagoes." The specific address is not given but in 2024, N. Dubuque St. runs from downtown Iowa City to today's Interstate 80, all within the city limits of Iowa City. |
|  | Iowa City Daily Republican (1876-1916) | "Some Observations on the Cicada", June 11, 1888: A copy of the same article from the Vidette-Reporter. |
|  | Iowa State Press (1866-1903) | "The Seventeen Year Locusts", H.F. Wickham, June 13th, 1888: Article anticipating the emergence of adult cicadas but that also notes the recent appearance of pre-emergence holes in great numbers in the area. |
|  | Slovan Americky (1873-1897) | May 23rd, 1888 (in Czech): anticipatory story about the coming emergence of 17-year cicadas. Notes that there are currently many waiting in the ground, especially under trees. |
| 1905 | Iowa City Daily Press (1904-1920) | 1) "Hated locust makes his bow" 17-year curse appears in Iowa City, May 19th, 1905: describes the discovery of cicada tunnels, pre-emergence, in a yard at the corner of Bloomington and Linn in downtown Iowa City: "Fully three hundred of the winged things were unearthed this morning by Will J. Lorack, in his garden, and brought down town for the inspection of scientists. Some of the onlookers call the insects grubs, only, but the university zoologists recognize the dreaded cicadas identity with the 17-year bane." 2) "Pickle in store for sneak thieves", June 20, 1905: this article about a spate of robberies in the city mentions the "...voice of the locust - the seventeen-year brand" being heard throughout the city. |
|  | The Iowa Citizen (1891-1907) | 1) "Locusts Due this Year", April 26, 1905: An anticipatory story about the expected emergence later in the summer, but refers to previous emergences in Johnson County: "I find that the periodical cicada or seventeen year locusts have appears in the summers of 1937, 1854, 1872, 1888...The swarm has much decreased in numbers since it was first noticed here." 2) No title, May 19, 1905: a brief note saying that cicadas "...are plentiful in the ground..." 3) "Sparrows have redeeming trait," June 14, 1905: Brief story about how English sparrows have been eating the 17-year cicadas. Confirms presence of cicadas in Iowa City: "...the sidewalks of the city are liberally strewn with the wings of the destructive insect and thanks are due to the sparrow." |
|  | Iowa City Citizen (1907-1916) | "Cicada is busy stinging trees", August 29th, 1911: Though clearly describing annual cicadas in the moment, is article refers to a mass emergence "...some seven or eight years ago...(probably six years, in 1905)...on the Kimball Road". This road connects with N. Dubuque St., mentioned as an emergence site in an 1888 article. |

Table S2. Classification accuracy of the 1930 and 2023 imagery.

|  | Overall | Kappa | Producer (Other) | Producer (Tree) | User (Other) | User (Tree) |
| --- | --- | --- | --- | --- | --- | --- |
| 1930 | 93.4% | 0.782 | 94.0% | 90.5% | 98.0% | 75.3% |
| 2023 | 92.2% | 0.768 | 91.9% | 94.6% | 98.4% | 72.5% |

**Table S3. Results of binary logistic regressions of 2023 *Magicicada* chorusing versus tree cover in 1930, 2023, 1836-1859 ("GLO") for 301 forested sites in Johnson County, IA.** Tree cover was calculated based on a circular area around each site of radii ranging from 0.5 km to 3.0 km. Also tested were "continuous" tree cover (present in both 1930 and 2023) and various combinations of tree cover and time periods considered together. Results include R2 values, AIC, and P-values.

|  |  | **R^2^** | | | | | | **AIC** | | | | | |
| --- | --- | --- | --- | --- | --- | --- | --- | --- | --- | --- | --- | --- | --- |
| **radius (km)** | **Area (km^2^)** | **1930** | **2023** | **Cont** | **GLO** | **1930+2023** | **19+23+GLO** | **1930** | **2023** | **Cont** | **1860** | **1930+2023** | **19+23+GLO** |
| **0.5** | 0.785398163 | 0.171 | 0.428 | 0.272 | 0.095 | 0.444 | 0.456 | 385.2 | 310.5 | 356.9 | 406.6 | 307 | 303.4 |
| **0.75** | 1.767145868 | 0.258 | 0.427 | 0.346 | 0.106 | 0.479 | 0.493 | 361.8 | 309.6 | 334.7 | 404 | 296.5 | 289 |
| **1** | 3.141592654 | 0.331 | 0.431 | 0.409 | 0.118 | 0.51 | 0.533 | 341.4 | 308.2 | 315.3 | 401.2 | 288.1 | 274.3 |
| **1.25** | 4.908738521 | 0.443 | 0.42 | 0.449 | 0.129 | 0.53 | 0.559 | 309 | 312 | 301.6 | 398.6 | 281.2 | 262.8 |
| **1.5** | 7.068583471 | 0.512 | 0.412 | 0.476 | 0.138 | 0.538 | 0.568 | 286.7 | 314.7 | 290.8 | 396.1 | 276.9 | 257.4 |
| **2** | 12.56637061 | 0.525 | 0.411 | 0.477 | 0.153 | 0.525 | 0.571 | 277.5 | 315.4 | 289.6 | 392.1 | 278.6 | 253.4 |
| **3** | 28.27433388 | 0.501 | 0.422 | 0.443 | 0.178 | 0.502 | 0.567 | 280.6 | 312.3 | 300 | 385.4 | 282.5 | 250.7 |

|  | **P-value** | | | | | |
| --- | --- | --- | --- | --- | --- | --- |
| **radius (km)** | **1930** | **2023** | **Cont** | **GLO** | **1930+2023** | **19+23+GLO** |
| **0.5** | <.0001 | <.0001 | <.0001 | <.0001 | .0253, <.0001 | .0249, <.0001, .0433 |
| **0.75** | <.0001 | <.0001 | <.0001 | <.0001 | <.0001, <.0001 | .0001, <.0001, .0280 |
| **1** | <.0001 | <.0001 | <.0001 | <.0001 | <.0001, <.0001 | <.0001, <.0001, .0036 |
| **1.25** | <.0001 | <.0001 | <.0001 | <.0001 | <.0001, <.0001 | <.0001, <.0001, .0010 |
| **1.5** | <.0001 | <.0001 | <.0001 | <.0001 | <.0001, .0025 | <.0001, <.0001, .0010 |
| **2** | <.0001 | <.0001 | <.0001 | <.0001 | <.0001, .7750 | <.0001, .0009, .0001 |
| **3** | <.0001 | <.0001 | <.0001 | <.0001 | <.0001, .5734 | .0015, .0029, <.0001 |

|  | **Z-score** | | | | | |
| --- | --- | --- | --- | --- | --- | --- |
| **radius (km)** | **1930** | **2023** | **Cont** | **GLO** | **1930+2023** | **19+23+GLO** |
| **0.5** | 5.68 | 8.33 | 6.98 | 4.13 | 2.24, 7.42 | 2.24, 7.12, -2.02 |
| **0.75** | 6.83 | 8.52 | 7.77 | 4.38 | 4.01, 7.18 | 3.86, 6.89, -2.2 |
| **1** | 7.6 | 8.52 | 8.18 | 4.64 | 4.87, 6.75 | 4.6, 6.57, -2.91 |
| **1.25** | 8.39 | 8.42 | 8.38 | 4.88 | 5.56, 5.14 | 5.18, 5.55, -3.28 |
| **1.5** | 8.7 | 8.35 | 8.67 | 5.09 | 5.81, 3.03 | 5.19, 4.33, -3.30 |
| **2** | 9.01 | 8.26 | 8.76 | 5.39 | 5.47, .29 | 4.38, 3.31, -3.88 |
| **3** | 8.86 | 8.19 | 8.7 | 5.84 | 4.69, -.56 | 3.17, 2.98, -4.51 |
